# Integrative analysis of CAKUT multi-omics data

**DOI:** 10.1101/2023.06.29.547015

**Authors:** Jumamurat R. Bayjanov, Cenna Doornbos, Ozan Ozisik, Woosub Shin, Núria Queralt-Rosinach, Daphne Wijnbergen, Jean-Sébastien Saulnier Blache, Joost P. Schanstra, José M. Fernández, Rajaram Kaliyaperumal, Anaïs Baudot, Peter A.C. ’t Hoen, Friederike Ehrhart

## Abstract

Congenital Anomalies of the Kidney and Urinary Tract (CAKUT) is the leading cause of childhood end-stage renal disease and a significant cause of chronic kidney disease in adults. Genetic and environmental factors are known to influence CAKUT development, but the currently known disease mechanism remains incomplete. Our goal is to identify affected pathways and networks in CAKUT, and thereby aid in getting a better understanding of its pathophysiology. Multi-omics experiments, including amniotic fluid miRNome, peptidome, and proteome analyses, can shed light on foetal kidney development in non-severe CAKUT patients compared to severe CAKUT cases. We performed FAIRification of these omics data sets to facilitate their integration with external data resources. Furthermore, we analysed and integrated the omics data sets using three different bioinformatics strategies. The three bioinformatics analyses provided complementary features, but all pointed towards an important role for collagen in CAKUT development. We published the three analysis strategies as containerized workflows. These workflows can be applied to other FAIR data sets and help gaining knowledge on other rare diseases.

## Introduction

Congenital Anomalies of the Kidney and Urinary Tract (CAKUT) cover a wide range of structural malformations that result from defects in the morphogenesis of the kidney and/or urinary tract ^1^. CAKUT affects three to six individuals per 1000 live births and constitutes the leading cause (∼40%) of end-stage renal diseases in childhood, and are significant contributors to chronic kidney disease in adults ^2,3^. In recent years, alterations in more than 50 genes have been shown to be associated with CAKUT, but a clear genotype-phenotype relationship remains absent ^3^. Therefore, in order to gain a better understanding of the disease, researchers should focus on molecular pathways and networks connecting genotype and phenotype. This requires multi-omics analysis and integration.

CAKUT is one of the rare diseases that was selected as a case study for the European Joint Programme on Rare Diseases (EJP RD). EJP RD focuses on integrating the fragmented raredisease research carried out in multiple countries. The objective is to facilitate data sharing and aggregation, and thereby enable higher statistical power. More precisely, the project aims to apply findable, accessible, interoperable, and reusable (FAIR) data principles ^4^ to the (meta)data and workflows, in order to allow a seamless data exchange between international scientific groups. Similarly, the software tools created during the course of the EJP RD project will be dispatched and applied on other rare disease data sets.

In this paper, our objectives are to illustrate with the CAKUT case study the benefit of multiomics integrative analysis and data FAIRification. We applied several bioinformatics strategies to a multi-omics data set from non-severe CAKUT patients and severe CAKUT patients. The main aims of our study were (1) to identify molecular pathways/networks that differentiate the severity of CAKUT conditions, and thereby contribute to the understanding of CAKUT molecular mechanisms; (2) to evaluate complementing features of different analysis methods for multi-omics data sets within the context of a rare disease. Furthermore, (3) we intend to make the data and the analyses FAIR and available for re-use. This supports open science in the rare disease field.

## Results

To increase our understanding of CAKUT disease aetiology, we performed multi-omics analyses on a total of 162 amniotic fluid samples. The omics types include previously published peptidome and proteome data from non-severe CAKUT and severe CAKUT patients, supplemented with a novel miRNome (See Methods section). We applied three different bioinformatics workflows to analyse and integrate this multi-omics data set. The workflows include intrinsic analysis using unsupervised (mixOmics) and supervised (momix) approaches, and extrinsic data analysis based on prior knowledge databases (pathway-level analysis). Each of the three complementary workflows used at least two types of omics data (Figure 1). In order to facilitate data integration and analysis, both data and analysis scripts were FAIRified.

**Figure 1.**
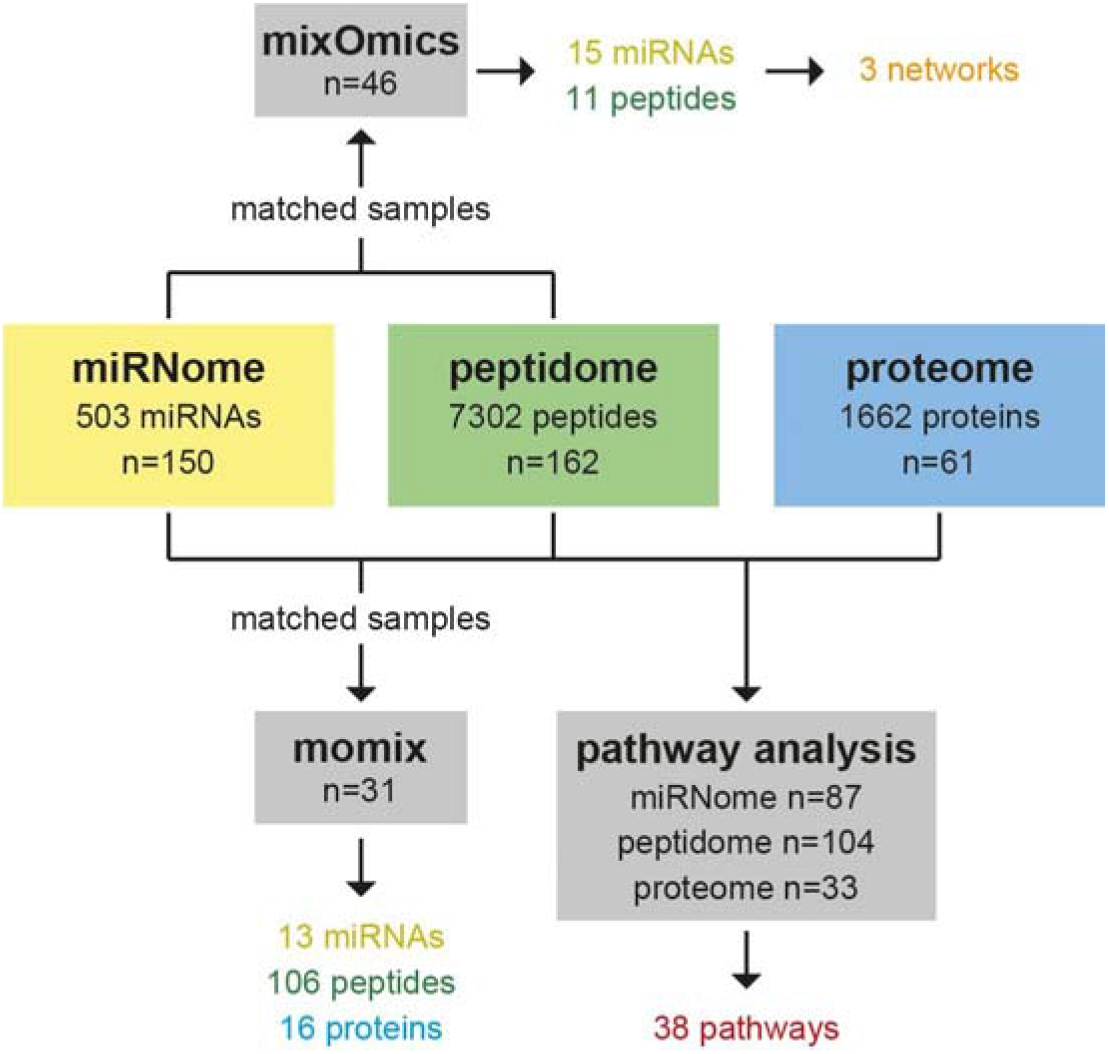
Analysis samples. Specification of the number of samples from each of the three omics data sets that was used for the three bioinformatics strategies (mixOmics, momix, and pathway analysis). For the mixOmics and the momix analysis, the number of samples was reduced, since these methods required samples from the same patient that matched between the different omics data sets. For the pathway analysis methods, all samples with sufficient clinical data were used for analysis. The results of each strategy are highlighted.

### Best place for Figure 1

#### FAIR data point creation

The omics data sets were FAIRified in order to promote findable, accessible, interoperable, and reusable data use. Furthermore, a new catalogue was created in the EJP RD FAIR Data Point (FDP), which was supplemented with the CAKUT data set descriptions [https://w3id.org/ejp-rd/fairdatapoints/wp13/catalog/4cad6f79-a7e1-46ef-8706-37f942f4aaea]. This promotes reproducibility and reusability of the data in future analyses.

#### Multi-omics integrative analysis with mixOmics

As a first approach to analyse the CAKUT multi-omics data, we used mixOmics, combining the miRNome and peptidome data with the mixOmics package ^5^. This approach identifies common patterns among multiple omics datasets by projecting data into a small number of dimensions, where the number of dimensions or components can be specified. Only the samples that matched between the two omics data sets and were in the training cohort of the peptidome study ^6^ were used (n=46; 30 non-severe and 16 severe CAKUT cases), due to the nature of the analytic approach in the supervised classification method of the mixOmics package. In the mixOmics analysis, the proteomics data were not used, because there were a limited number of matching samples compared to the miRNome and peptidome data (Figure 1).

As part of the mixOmics analysis, Partial Least-Squares Discriminant Analysis (PLS-DA) and sparse PLS-DA (sPLS-DA) were used to identify a subset of variables that could explain the variance between non-severe CAKUT and severe CAKUT patients. It was noted that the peptidome data has a higher variance than miRNA data for the first two components in both PLS-DA and sPLS-DA analyses ^7^ (Table 1 and Figure 2A), which indicates that the peptidome data has a better segregation of non-severe CAKUT versus severe CAKUT patients then the miRNome. The main variance between the groups emerged from peptides that were derived from a variety of collagen proteins (Figure 2B). These observations confirmed the findings obtained using the peptidome data alone ^6^. Although classification accuracy is higher when the peptidome data alone was used, multi-omics analysis revealed relationships between the miRNome and the peptidome (Figure 2C). These mainly include positive correlations for a large number of miRNAs with only three peptides. Only one negative relation was observed between one of the COL1A1 peptides and mir-hsa-6768-5p. For the highest scoring miRNAs and peptides of the mixOmics analysis, a network-based visualisation was performed. This network-based visualisation revealed a large collagen and cytoskeleton cluster (Figure 2C). Furthermore, unsupervised analysis, shown by heatmap clustering (Figure 2D), confirmed strong correlations between certain peptides and miRNAs. In conclusion, the mixOmics method indicates an important role for collagen on miRNome and peptidome level.

**Table 1.** Contribution scores per omics data for each of the two principal components of the principal component analysis, where sPLS-DA, a variable selection method was applied to select the optimal number of peptides and miRNAs.

| Method | Component 1<br>(miRNA/peptidome) | Component 2<br>(miRNA/peptidome) |
| --- | --- | --- |
| PLS-DA | 80.47% / 96.70% | 80.77% / 98.35% |
| Sparse PLS-DA | 79.40% / 96.43% | 85.44% / 96.15% |

**Figure 2.**
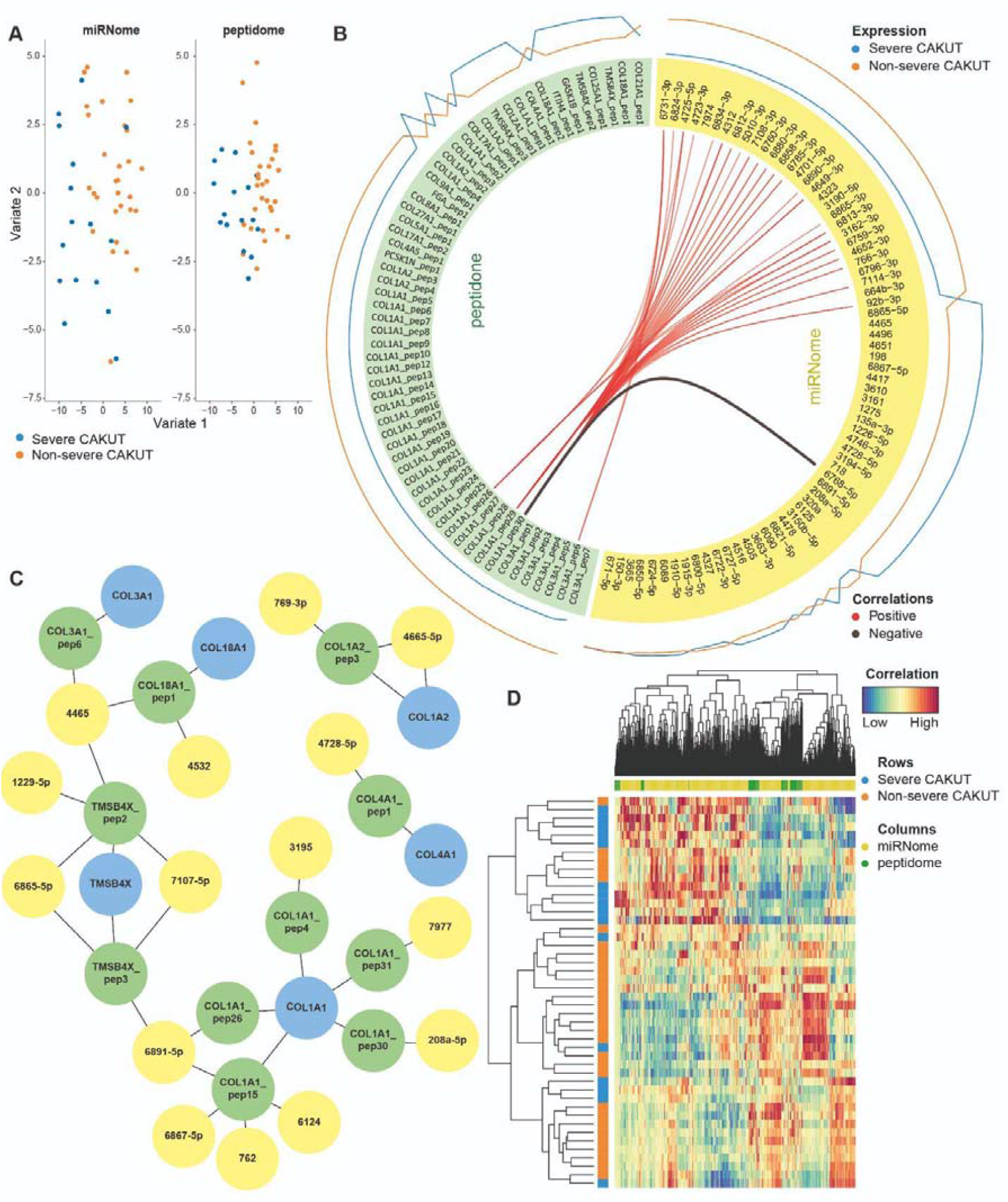
Integrative analysis of miRNome and peptidome data to identify combinations of variables from both omics datasets in comparison to the single-omics analysis using only peptidome data. Figure 2A: Multi-omics integration of miRNome and peptidome data using the block sPLS-DA method of the mixOmics package. Peptidome and miRNome data were matched by patient. Variates 1 and 2 indicate different latent components, where both peptidome and miRNome data are projected onto a smaller 5-dimensional subspace (see Methods). Figure 2B: Circos plot of correlations based on the sPLS-DA results using the miRNome (yellow) and peptidome (green) data for the first two components. miRNAs are indicated by their hsa-miR identifiers. Peptides are mapped to their respective proteins and multiple matches to the same protein are shown with the numbered suffixes. The exact peptide sequences can be found in the Supplemental Table 1. Only correlations scoring above 0.80 are shown. Figure 2C: Network-based integration of the miRNome, peptidome, and proteome data sets to depict the most relevant molecules identified by the mixOmics approach. The network is composed of the most relevant miRNAs (yellow) and peptides (green) based on sPLS-DA analysis as described in the Methods section. In this case, the peptide sequences were used to map peptides to proteins (blue) using sequence alignment (see Methods). Peptides and miRNAs are indicated as in 2B. The larger network is a collagen and cytoskeleton network consisting of COL3A1, COL18A1, TMSB4X involved in cytoskeleton organisation, and COL1A1. The two smaller networks also include COL1A2 and COL4A1. Figure 2D: Unsupervised analysis between miRNAs and peptides displayed by a heatmap. The colours are based on their contributions to the first two components. Only miRNA and peptides with correlations above 0.80 are shown.

### Best place for Table 1 and Figure 2

#### Joint multi-omics dimensionality reduction analysis

In the second strategy, we applied eight different unsupervised joint dimensionality reduction methods on the peptidome, proteome, and miRNome data using the momix notebook ^8^. We used the 31 samples (18 non-severe CAKUT cases and 13 severe CAKUT cases) that matched between the three omics data sets. A joint dimensionality reduction method decomposes the omics datasets into omics-specific weight matrices and a joint factor matrix. We ran the dimensionality reduction methods to obtain the two most important factors (k=2). Most non-severe and severe CAKUT patients could be separated by one of these two factors, which segregate the two groups (Figure 3A-C). To evaluate the methods and choose the most relevant factor, we measured how well the two sample groups could be clustered. For each method and each factor, we used k-means clustering. We ran k-means 1000 times and counted the number of samples that were in the correct cluster in accordance with the clinical diagnosis. The baseline accuracy is 58% (18 over 31); it can be obtained by assigning all the samples to one of the two clusters. The accuracies of the joint dimensionality reduction methods range from 65% to 90% when from the two factors the better segregating one is taken into account (Table 2).

**Figure 3.**
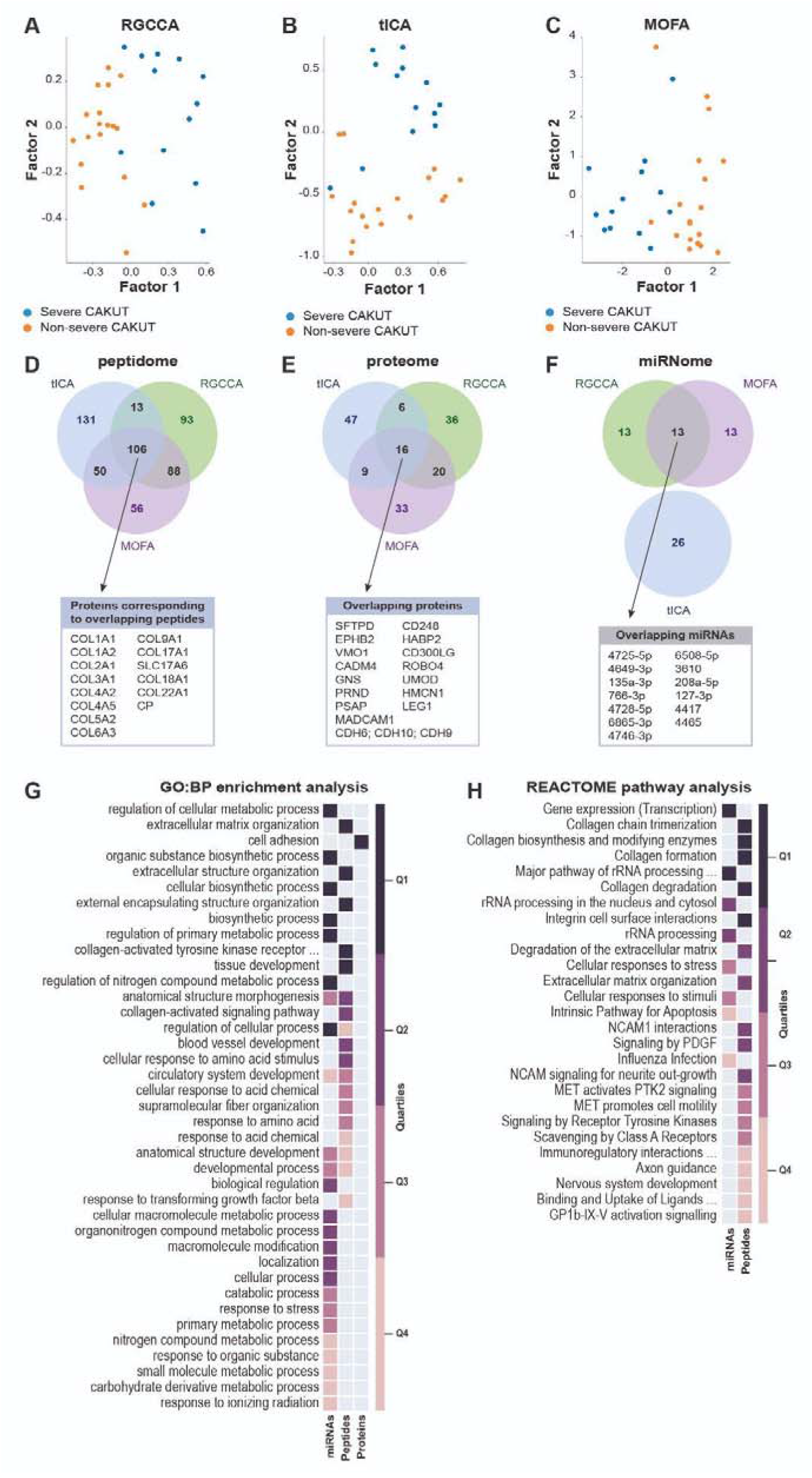
Joint multi-omics dimensionality reduction analysis. Figure 3A-C: Projections of all samples on the first two factors obtained by (a) RGCCA, (b) tICA and (c) MOFA. Figure 3D-F: Overlap of top 5% peptides, proteins, and miRNAs selected by RGCCA, tICA, and MOFA analysis. Figure 3G: GO Biological Process enrichment analysis results of the features selected from different omics data by multiple methods. The significant results from different omics are filtered and integrated by orsum. The colours indicate the quartile of the rank of the significant term for the specific dataset. Enrichment scores can be found in Supplementary Table 5. Figure 3H: Reactome enrichment analysis results of the features selected from different omics data by multiple methods (there is no enrichment result for genes selected from the proteome data). The significant results from different omics are filtered and integrated by orsum. The colours indicate the quartile of the rank of the significant term for the specific dataset.

**Table 2.**
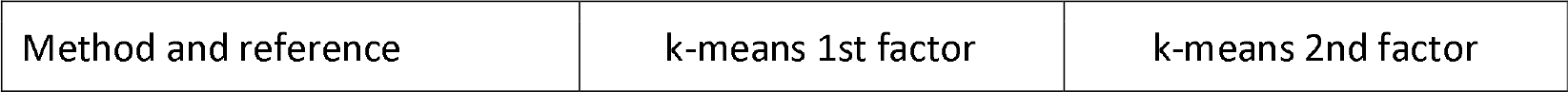
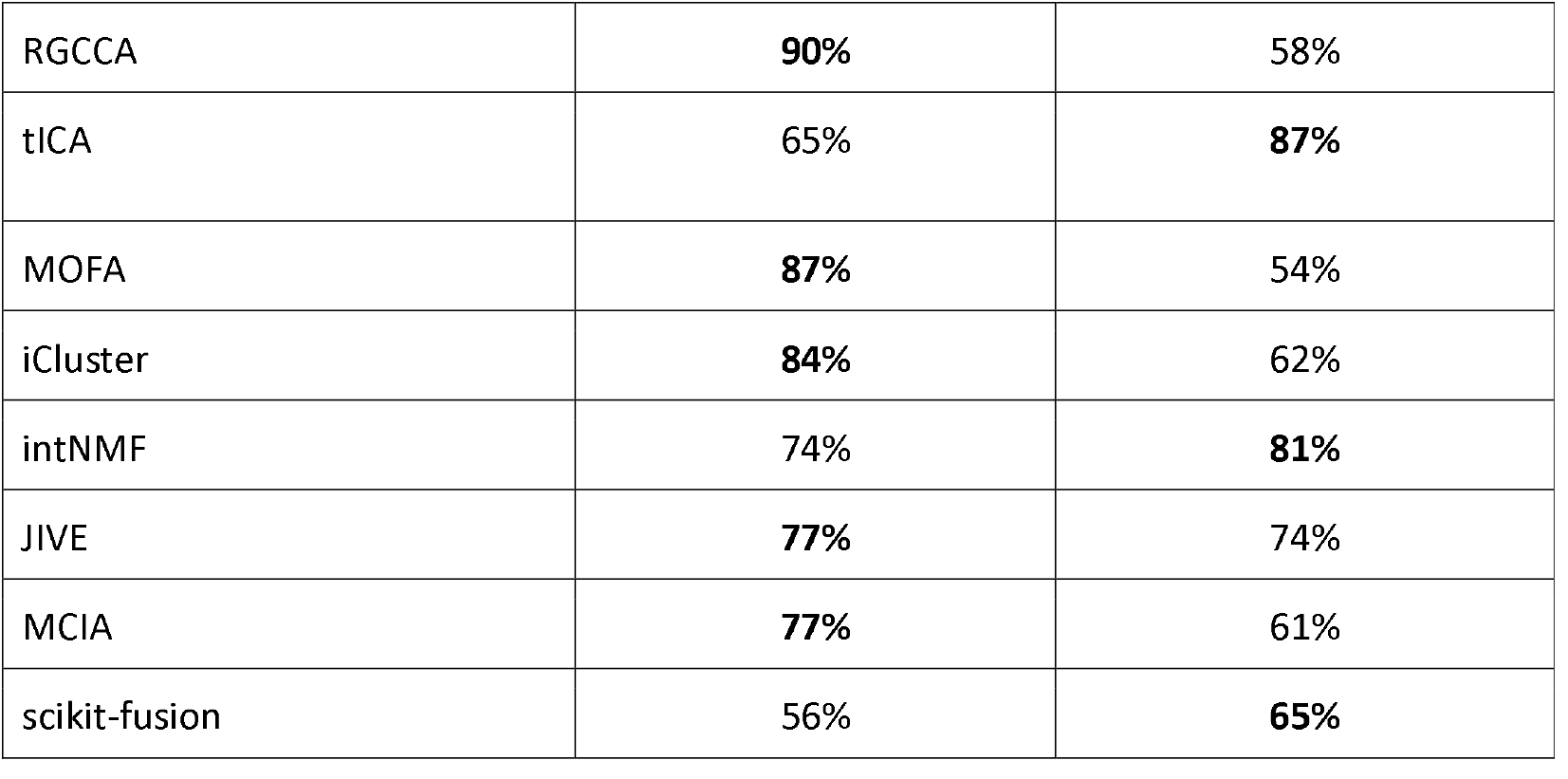
Accuracy of k-means clustering runs on each one of the two factors calculated by joint multi-omics dimensionality reduction methods. The bold numbers represent the higher accuracy obtained by each method.

| Method and reference | k-means 1st factor | k-means 2nd factor |
| --- | --- | --- |
| RGCCA | **90%** | 58% |
| tICA | 65% | **87%** |
| MOFA | **87%** | 54% |
| iCluster | **84%** | 62% |
| intNMF | 74% | **81%** |
| JIVE | **77%** | 74% |
| MCIA | **77%** | 61% |
| scikit-fusion | 56% | **65%** |

### Best place for Table 2

Based on the accuracy, we selected the three methods that were the most successful in separating non-severe CAKUT patients and severe CAKUT patients, namely RGCCA, tICA, and MOFA (Figure 3A-C). Within the weight matrices created by these methods, we used the weight vectors corresponding to the better of the two factors. We then used the absolute value of the weights assigned to the features and selected the top 5% of peptides, proteins, and miRNAs from each method for further analysis (Supplementary Table 2-4).

### Best place for Figure 3

We focused on the peptides and proteins identified as top 5% by all three methods, and miRNAs identified by two methods, as there was no miRNA in common to all three methods (Figure 3D-F). This resulted in 106 peptides, 16 proteins, and 13 miRNAs. The 106 peptides correspond to 15 proteins, mainly collagens. None of these 15 proteins were identified in the top 5% of the proteome. However, some corresponded to related proteins. For instance, Cadherins (CDH6, CDH9, CDH109) and CADM4 play a role in calcium-dependent cell adhesion. Furthermore, ROBO4, UMOD, HABP2, MADCAM1, HMCN1, and EPHB2 have been indicated to be involved in cell adhesion, cell junctions and/ or the migration of one or more specific cell types. Overall, this indicates that the peptidome and the proteome identify different proteins but similar processes.

We performed enrichment analysis to identify the most important biological processes associated with the selected peptides, proteins, and miRNAs (Supplementary Table 2-4). In this analysis, we used the proteins selected from the proteome data, the proteins corresponding to the selected peptides and the genes targeted by the selected miRNAs. We used orsum ^9^ in order to present the enrichment results and to filter redundant annotation terms (Figure 3G). Five Gene Ontology Biological Process (GO-BP) terms are significantly enriched in both the miRNome and peptidome data, mainly indicating misregulation of organ structure and development in non-severe CAKUT patients versus severe CAKUT patients. “Cell Adhesion” is the only significantly enriched GO-BP term in the proteomics data. Proteins corresponding to the selected peptides are further enriched in extracellular processes, including the process entitled “collagen-activated tyrosine kinase receptor signaling pathway”. Cell adhesion and collagen related pathways are also significant when REACTOME pathways are used in the enrichment analysis (Figure 3H). Finally, for the miRNome data, the REACTOME enrichment analysis of the genes targeted by the selected miRNAs mainly revealed rRNA and transcription processes. Deregulations of these processes are, to the best of our knowledge, not described in CAKUT. The GO-BP enrichments indicated a role for the miRNA regulated genes in metabolomics and biosynthesis for which misregulation could affect organ structure and development.

### Pathway-level analysis

We analysed the CAKUT omics data for overrepresented pathways within WikiPathways database ^10^. From 634 pathways in the database, 38 pathways were overrepresented and had a link between miRNA and protein (or peptide mapped into protein) based on the CAKUT patient data. In these pathways, we found 15 links between miRNome and proteome where both interaction partners are significantly differentially expressed. The PI3K-Akt Signalling Pathway (WikiPathways:WP4172) ^11^ contained five links between miRNAs and the peptidome or proteome (Figure 4A). The 10 remaining links between miRNA and proteins are indicated in Figure 4B.

**Figure 4.**
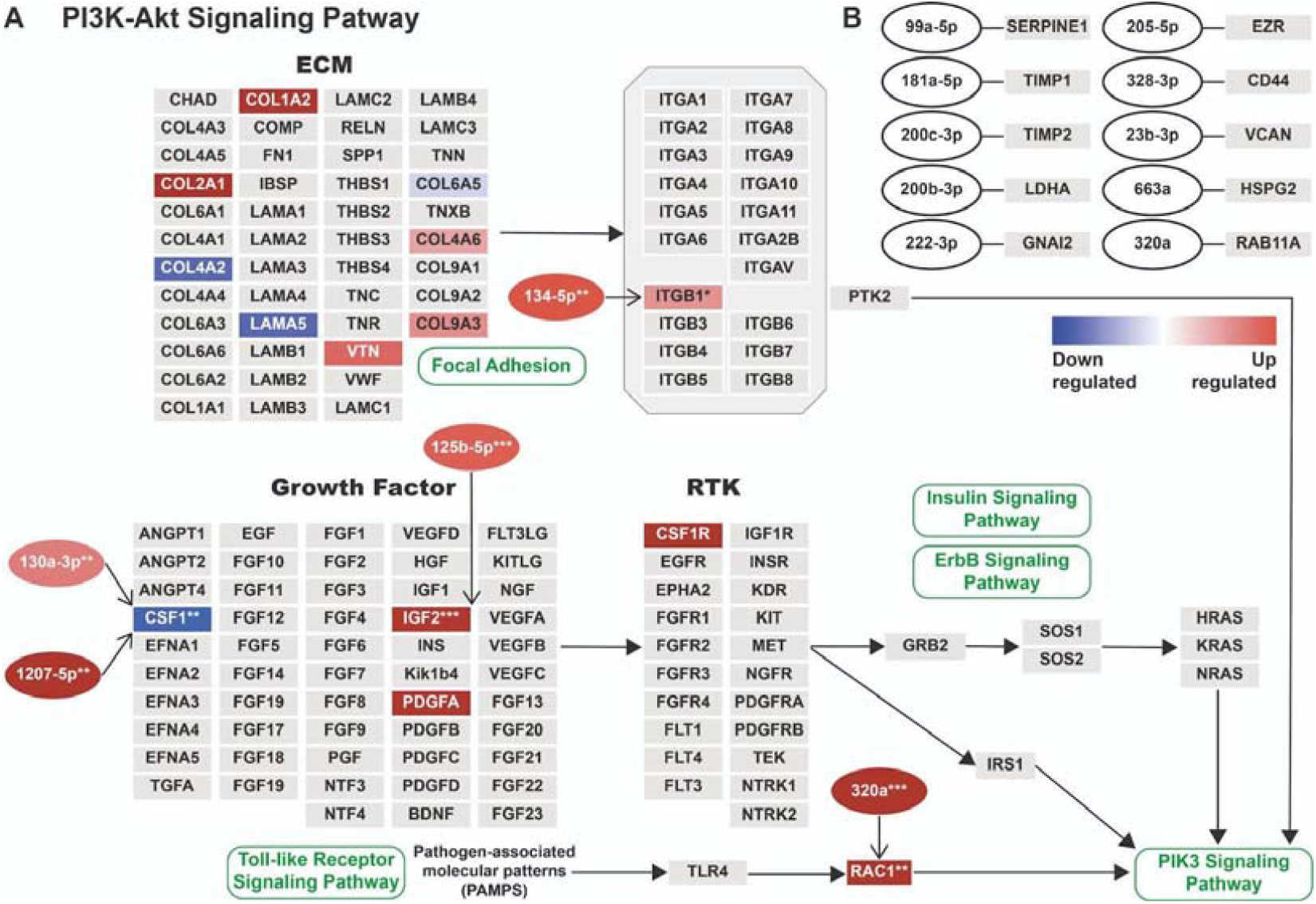
Pathway enrichment analysis. Visualisation of the interacting differentially expressed proteins/peptides/miRNAs in WikiPathways pathway database on the combined miRnome, peptidome, and proteome data. Rectangular nodes represent protein products as determined from the peptidome and proteome, ellipses represent miRNAs indicated by their hsa-miR identifiers. A) Visualization of the PI3K-Akt Signalling Pathway as adjusted from WikiPathways (WikiPathways:WP4172). Only a part of the pathway is shown from the larger pathway to emphasise the section where most differential expressions occurred. Blue indicates downregulation and red upregulation, as indicated by the gradient bar. Asterisks indicate the enrichment significance (p-value). Grey nodes mean that the expression data did not exist or did not have significant differential expression (p<0.05). On the one hand, we found that IGF2, ITGB1, and RAC1 were upregulated in the same direction as their miRNAs. On the other hand, CSF1 was downregulated in contrast to its targeting miRNAs, which were both upregulated. B) The 10 remaining significantly linked miRNA and proteins, from the 15 interactions that were identified in total. The gene products were selected only when either peptidome or proteome indicated significant levels of differential regulation, as well as, the significant miRNAs targeting them.

The PI3K-Akt pathway also includes certain collagens that had been associated with CAKUT in the original study ^6^. However, we could not identify any significant links between these proteins and the miRNome. Instead, in this pathway, we found significant links between four gene products (CSF1, IGF2, ITGB1, and RAC1) and five miRNAs (hsa-miR-130a-3p, hsa-miR- 1207-5p, hsa-miR-125b-5p, hsa-miR-134-5p, and hsa-miR-320a).

### Best place for Figure 4

## Discussion

Using the output from the **mixOmics** approach a network was identified related to collagen and cytoskeleton remodelling, consisting of COL3A1, COL18A1, TMSB4X, and COL1A1, and two smaller networks including COL1A2 and COL4A1. In this analysis, we used a supervised classification approach with the mixOmics method of the mixOmics package, which requires matching samples among omics data sets. Since the number of overlapping samples in all sets decreased when the proteomics data were included in the mixOmics-based analysis, we decided to exclude the proteomics data for this specific analysis. MixOmics proposes two approaches, sPLD-DA and PLD-DA. The difference between sPLS-DA and PLS-DA was insignificant, probably because sPLS-DA is expected to be beneficial over PLS-DA for high dimensional data^12^. Additionally, mixOmics analysis allowed the identification of a collagen-related cluster solely based on the peptidome and miRNome data. The main variance in the data stemmed from a range of miRNAs that could be connected to a handful of peptides (Figure 2B). Most of these relations were positive correlations, while only hsa-miR-6768-5p and COL1A1_pep30 (ADGQpGAKGEpGDAGAKGDAGPpGP) had a negative correlation. hsa-miR-6768-5p has not been previously identified or predicted to affect COL1A1. While an important role for collagen in CAKUT was previously established ^6,13,14^, COL1A1 has not specifically been linked to CAKUT. Furthermore, this work highlights potential novel miRNA and peptide relations, which might be most relevant to study to get a better understanding of CAKUT.

**Unsupervised joint dimensionality reduction analysis with the momix notebook** identified the most relevant molecules from the three omics data sets. We further selected the results of the three best performing joint dimensionality reduction methods among the eight tested. From the proteome analysis, CDH6 (P55285), CDH9 (Q9ULB4), and CDH10 (Q9Y6N8) are particularly interesting, as these cadherins regulate hippo signalling, which plays a role in kidney and urinary tract development (Figure 3E) ^15,16^. Furthermore, UMOD (P07911) was previously associated with medullary cystic kidney disease, familial juvenile hyperuricemic nephropathy, and glomerulocystic kidney disease ^17^. Whether mutations in UMOD are a cause of CAKUT is still under debate ^17^. The peptide analysis revealed COL4A5 (P29400) as an interesting protein, as it is one of the glomerular basement membrane proteins that cause Alport syndrome ^18^. COL4A1 (P02462) is also of interest. This protein is identified by all the three best performing methods due to different peptides (MOFA and RGCCA found the peptide COL4A1_pep1, tICA found the peptide COL4A1_pep2) (Supplemental Table 1) and it is associated with kidney diseases ^13,14,19^.

Comparing the momix and mixOmics workflows, there is an overlap in the identified molecules of interest, including COL1A1, COL1A2, COL3A1, and COL18A1. In the GO enrichment analysis, we obtained different annotation terms indicating that momix and mixOmics approaches are complementary.

The **analysis at pathway-level** used the molecular interactions of WikiPathways, a pathway database extended with miRNA-target information as a backbone to investigate the interactions of interest. The advantage of this method is that it integrates prior knowledge into the analysis, which is especially important when the signal extracted from the data is low. Using this pathway analysis method, we identified 15 functional links between significant differentially expressed proteins and the miRNome. The PI3K-AKT signalling pathway hosts five of these interactions between the different omics data sets, making this the most relevant pathway for CAKUT disease progression (Figure 4A). In addition, it harbours several collagen proteins previously identified by the other methods as well. A role for collagen in CAKUT disease development was previously established ^6,14,20^. The other interactions from i.a. Focal Adhesion (WikiPathways:WP306) or Senescence and Autophagy (WikiPathways:WP28806) pathway, are shown in Figure 4B. The limitation of the pathway-level analysis is the dependence on knowledge databases of molecular interactions. Nonetheless, for both pathways and miRNA-target interactions, there are several options regarding analysis. On the one hand, WikiPathways is an open, community created, and expert curated database ^10^. The contributions that define the content are dependent on published literature, and the pathways undergo regular curation to be updated with current findings. On the other hand, miRTarBase is a miRNA-target interaction database that provides manually selected, experimentally validated miRNA-target interactions from published literature ^21^. Integrating analysis methods using these and other databases to cross validate the information measured on patients relevant conclusions for disease research.

Altogether, the different bioinformatics strategies and methods presented in this paper offer a complementary spectrum of possible multi-omics analyses, the analysis of rare disease data sets. Notably, most of these methods identified the same (functional) group of genes, with differences on the weighing of correlation statistics or the use of prior knowledge supported methods. Importantly, methods based on mathematical analysis, ignoring existing biomedical knowledge, allow us to identify potentially interesting features in a hypothesis free manner. Pathways, or approaches based on prior knowledge in general allow us to select functional and molecular interactions from the given data to support a biomedical interpretation of the results. A combination of these strategies is expected to be advantageous for the analysis of (multi)omics data in the field of rare diseases.

There is an increasing demand towards open science, which requires making the **data, analysis tools, and whole workflows FAIR**ly available together with the results. This significantly increases the possibility to reproduce results and counteract the current crisis in reproducibility and trust in scientific studies. This demand is especially high in the rare disease field where the naturally limited number of patients, samples, and data has ever since encouraged international and interdisciplinary collaborations to pool data and exchange methods in how to deal with low sample numbers. To this purpose, we hope to aid research reliability and reproducibility by providing both FAIR metadata and workflows as presented in this study and supported by the EJP RD.

## Conclusion

With this study, we provide several different bioinformatics strategies that can identify biologically relevant biological molecules, pathways, and networks from multi-omics data sets in an unsupervised and supervised manner. The identified proteins, peptides, and miRNAs highlight modules relevant for CAKUT disease and can be used for future investigations. Finally, the application of open science principles in this study contributes to the reusability of data and workflows in, but not limited to, the rare disease field.

## Methods

### Multi-omics data sets

The CAKUT multi-omics data set was obtained from a previously published study and reinvestigated in collaboration with the authors of the original study ^6,22^. The initial study contains amniotic fluid samples from proteome and peptidome. Here we added novel miRNome data from amniotic fluid samples from the same patients as described below. In total 175 individuals were studied, of which 104 samples had a clinical diagnosis. This diagnosis consisted of antenatal diagnosis, amniotic fluid phenotype, and the postnatal outcome at two years of age. Non-severe CAKUT patients had normal GFR (glomerular filtration rate), moderately reduced GFR (60 to 90 ml/min per 1.73 m^2^), or reduced GFR (<60 ml/min per 1.73 m^2^) postnatal outcome, but are marked as CAKUT patients based on their antenatal diagnosis. The severe CAKUT patients were diagnosed with severe renal failure, end-stage renal disease, or the renal phenotype lead to a termination of pregnancy.

In total, the abundances of 7302 peptides were measured in the amniotic fluid samples of 162 subjects, 503 miRNAs were significantly detected from 150 samples and 1662 proteins were detected in 61 samples. Each of these three omics data sets includes non-severe CAKUT vs. severe CAKUT cases.

## miRNome sample collection and analysis

For miRNA analysis, amniotic fluid samples were collected in a prospective multicenter observational study focusing on foetal bilateral CAKUT as part of a clinical trial (https://clinicaltrials.gov/ct2/show/NCT02675686). The CAKUT disease severity was defined based on the renal status after two years of postnatal clinical follow-up.

The total RNA was isolated using the Agilent RNA 6000 Pico kit protocol (5067-1513) and microRNAs were profiled using Agilent microRNA slides (Sanger miRBase release 21). The samples were labelled and hybridised according to the Agilent’s microRNA Complete Labeling and Hybridization Kit protocol (5190-0456), followed by Spike-ins with the Agilent’s microRNA Spike-In Kit protocol (5190-1934) and analysed using Aligent’s High-Resolution Microarray Scanner GS2505_C. Features were called using the Agilent Feature Extraction software (version 11.0.1.1) and sample intensities were normalised using quantile normalisation (RMA). The miRNome data is available at https://zenodo.org/record/7866785#.ZE9wZXZBwuV.

### FAIR Data Point and data deposition

We used FAIR Data Point (FDP) to describe the CAKUT multi-omics data set. FDP is a metadata service that provides descriptions about resources ^23^. FDP uses Data Catalogue Vocabulary (DCAT) to capture the resources metadata. FDP serves descriptions of resources to both humans and machines, which makes integration of different data sets easier and allows reproducing results from previous studies. The human users who visit the FDP see the resource descriptions as HTML documents and the machine gets resource descriptions as a RDF document. We used the FDP [https://w3id.org/ejp-rd/fairdatapoints/wp13] created for the EJP RD project to describe CAKUT multi-omics data sets.

### Workflow specifications

Workflows from each of the three different types of analyses have been registered at the WorkflowHub [https://workflowhub.eu/projects/40], a scientific FAIR workflow registry. From this registry, each workflow can be downloaded together with all required scripts and data files as a single package ^24^. This facilitates re-analyses of the same data sets but also application of the workflows to additional data sets.

### Multi-omics integrative analysis with mixOmics

We used the mixOmics package ^25^ (version 6.10.9) with PLS-DA (Partial Least Squares Discriminant Analysis Discriminant Analysis) and sPLS-DA (sparse PLS-DA) supervised classification. The sPLS-DA method allowed for variable selection on each omics. As in principal component analysis, sPLS-DA projects large input data into a smaller dimensional space, with each component representing a different dimension. From the peptidome data, 53 samples were used for training and 51 samples were used for validation. The samples were assigned to each of the groups in the initial study ^6^. In line with this grouping, of the samples that matched between the peptidome and miRNome, 41 were used for training and 46 for validation. Nonetheless, when matching samples of the proteome to the other two omics data sets, there were only 23 and 10 samples for training and validation remaining respectively. Thus, the proteome data was not used in the mixOmics analysis. For the sPLS-DA classification, a maximum of five components were chosen, where each component represents a separate dimensional subspace for data projection. The first component uses 50 miRNAs and peptides, the second component uses 20 miRNAs and 10 peptides, whereas all other three components use all miRNAs and peptides. Peptides are mapped to their respective proteins and multiple matches to the same protein are shown with the numbered suffixes. The exact peptide sequences can be found in the Supplemental Table 1.

For the network shown in Figure 2C: (i) the edges between miRNAs and peptides are based on their statistical significance, which was based on the multi-omics analysis of the training data. (ii) The edges between miRNAs and proteins are identified based on the known miRNA-mRNA biological links provided by the mirTarBase database, version 8.0 ^21^. (iii) The edges between proteins and peptides are identified by aligning peptide sequences against protein sequences using NCBI BLAST.

We used R version 4.0.3 for the analysis. All R packages necessary to run these scripts are specified in the Docker file included at [https://workflowhub.eu/projects/40].

### Multi-omics integrative analysis with joint dimensionality reduction using momix

We applied eight joint dimensionality reduction methods on peptidome, proteome, and miRNome data for 31 samples (18 non-severe CAKUT cases and 13 severe CAKUT cases) matched in the different omics data sets. We used the momix notebook ^8^. The methods we used are iCluster ^26^, Integrative NMF (intNMF) ^27^, Joint and Individual Variation Explained (JIVE) ^28^, Multiple Co-Inertia Analysis (MCIA) ^29^, Multi-Omics Factor Analysis (MOFA) ^30^, Regularized Generalized Canonical Correlation Analysis (RGCCA) ^31^, matrix-tri-factorization (scikit-fusion) ^32^, and tensorial Independent Component Analysis (tICA) ^33^.

We ran the methods to obtain two factors in the reduced dimension. We observed that non-severe CAKUT and severe CAKUT patients could be separated by one of the two factors (Figure 3A-C). We used k-means clustering to select the factor better segregating non-severe CAKUT and severe CAKUT patients. We ran 1000 k-means clustering on each factor independently, and calculated the accuracy. The two labels are assigned randomly to the two clusters, the accuracy is measured; then labels are switched, accuracy is measured again; the higher accuracy allows selecting the better factor. Based on these accuracies, we also selected the best performing three methods. The top three methods, RGCCA, tICA and MOFA, obtained 87% or 90% accuracy using a single factor (Table 2). We used the results from these methods for subsequent analyses.

From the weight matrices created by the top three methods, we used the weight vectors corresponding to the selected better segregating factor. Using the absolute value of the weights assigned to the features, we selected the top 5% peptides, proteins and miRNAs. Among these top 5% molecules identified by the top three methods, we focused on the peptides and proteins identified by all three methods, and the miRNAs identified by two methods (RGCCA and MOFA), as there was no miRNA in common to all three methods.

For robustness, we used the top features identified by multiple methods. We used the peptides and proteins common to three methods, and miRNAs common to two methods, as tICA does not have any miRNAs common with the other two methods. This resulted in 106 peptides, 18 proteins, and 13 miRNAs (note that for proteins, 16 features are selected but one feature is “P55285;Q9Y6N8;Q9ULB4”, which correspond to three cadherins, CHD6, CHD9 and CHD10). The peptides were mapped to protein/gene using the UniProt Retrieve/ID mapping module (https://www.uniprot.org/uploadlists/) ^34^. For miRNA to target gene mapping, we used hsa_MTI.xlsx from miRTarBase (Release 8.0) ^21^.

For the enrichment analysis, we used g:Profiler ^35^. In the enrichment analysis, we used only CDH6 from the cadherins that were measured jointly in proteomics analysis (P55285; Q9Y6N8; Q9ULB4), to prevent inflation in the enrichment results. For the presentation and the filtering of redundant annotation terms in the enrichment results, we used orsum ^9^.

### Pathway-level analysis to detect functional links

The peptidome and the proteome data sets were quantile normalised and log2 transformed as previously described ^36,37^. Before transformation, peptide IDs were mapped to protein IDs, and were summarised into single protein-level values using geometric mean ^22^. The miRNome data set was already normalised and transformed, thus the information of their target genes could be added to each miRNA ID without additional data manipulation, using the information provided by miTaRBase. As a result, all three data sets had been mapped to their appropriate gene product-level identifiers.

Once the data sets were prepared, we applied one-predictor logistic regression for each protein-or miRNA-level and obtained the effect size (log2 fold change) and p-values. Each element of each data set (miRNA/peptide/protein) was deemed significantly differentially expressed if the corresponding p-value was less than 0.05.

In this analysis, we created an extended pathway network, using the WikiPathways repository (Version 20210110). For the pathway-level analysis, first each of the three omics data sets was analysed to identify overrepresented pathways. Subsequently, pathways associated with the significant miRNA-protein links were detected. A miRNA-protein link may possibly be implying causality, if both a miRNA and its target are differentially expressed. Pathways, which are overrepresented and contain at least one link from a significant miRNA either to a significant peptide or to a significant protein were identified. More specifically, a pathway was selected if it meets two conditions: (1) a gene product in the pathway was significantly differentially expressed by either the peptidome or proteome, or (2) there exists a miRNA, which targets the gene product, and the miRNA is significantly differentially expressed.

Finally, since the selected pathways only included information of gene products, they were extended using the miRNA targeting information when necessary. A visualisation of the selected pathway with study data and additional information was created.

## Data Availability

The CAKUT multi-omics data sets for the proteomics and peptidomics data sets are available with the original studies ^6,22^ and the repositories mentioned there. The miRNome data is available at https://zenodo.org/record/7866785#.ZE9wZXZBwuV. We added a FAIR data point to describe the CAKUT data sets here: <https://w3id.org/ejp-rd/fairdatapoints/wp13/catalog/4cad6f79-a7e1-46ef-8706-37f942f4aaea>.

## Code Availability

All data analysis codes are available at Workflowhub in this registry: https://workflowhub.eu/projects/40.

## Acknowledgements

The authors would like to thank Matthias Haimel, Laura Rodriguez-Navas, Franz Schäfer, Nazli Sila Kara, Ugur Sezerman, Daphne Wijnbergen, Ana Rath, and Chris Evelo for helpful discussions, and the anDREa team for technical support.

This work is supported by the funding from the European Union’s Horizon 2020 research and innovation programme under the EJP RD COFUND-EJP N° 825575. OOs work is supported by the Excellence Initiative of Aix-Marseille University - A*Midex, a French “Investissements d’Avenir” programme. JS and JBs part of the work was financed by the French “Programme Hospitalier de Recherche Clinique” (PHRC) number N° 10 138 01 - N° RCB 2010-AO1151-38, by grants from the “Fondation pour la Recherche Médicale” (grant number DEQ20170336759), and from the European Commission Seventh Framework Programme (FP7/2007-2013) grant agreement no. 305608 (EURenOmics).

## Author contributions

JSB and JS collected the data, JB, CD, DW, NQR, and RK did (FAIR) data management, data analysis was done by JB, OO, WS, CD, and FE, workflows and containerization by JB, CD, and JF. The first draft of the paper was written by JB, OO, WS, CD, AB, and FE, the study design was done and supervised by JB, PH, and FE. All authors critically reviewed the first draft, contributed writing the second draft, and have read and approved the final version of the paper.

